# Immunomodulatory Biomaterials Enhancing Implant Osseointegration: Knowledge Mapping of Research Evolution from March 2005 to March 2025

**DOI:** 10.1101/2025.05.29.656932

**Authors:** Li Yue, Liu Geng

**Affiliations:** Tianjin Stomatological Hospital, Nankai University Stomatological Hospital, Tianjin Key Laboratory of Oral Functional Reconstruction, Tianjin 300041, China

**Author notes:** Corresponding author : Li Yue.

**Keywords:** Immune regulation, Biomaterials, Implants, Osseointegration, Bibliometrics

## Abstract

**Objective:** To delineate the evolutionary trajectory of immunomodulatory biomaterials in implant osseointegration through bibliometric analysis, identifying pivotal theoretical breakthroughs and technological advancements.

**Methods:** A total of 419 articles (2005–2025) from the Web of Science Core Collection were analyzed using a multi-tool framework. Current research status and hotspots were evaluated by co-occurrence analysis of keywords and institutions using VOSviewer. The evolution and bursts of the knowledge base were assessed through co-citation analysis of references, authors, and journals via CiteSpace. Thematic evolution and keyword trends were mapped using the bibliometrix package in R.

**Results:** The field exhibited “intermittent-explosive” growth (32.7% annual increment), with China leading global contributions (69.4%). The osteoimmunomodulation (OIM) theory emerged as the cornerstone, emphasizing spatiotemporal macrophage polarization (M1/M2 balance) and multi-signal crosstalk (BMP-2/VEGF/OSM). Key technological pathways included: ① Surface engineering (nanotopography, ion-doped coatings); ② Smart materials (3D-printed scaffolds, pH/ROS-responsive carriers); ③ Antibacterial-immunomodulatory synergy. Burst detection revealed shifting frontiers toward clinical translation (2023-2025 burst: “3D printing”, strength=4.05) and precision modulation (“macrophage polarization”, strength=9.02).

**Conclusion:** Immunomodulatory biomaterials are transitioning from mechanistic exploration to clinical adaptation. Future development requires integrating dynamic microenvironment-responsive designs with multi-omics validation to address macrophage heterogeneity, ultimately enabling personalized osseointegration therapies.

## 1. Introduction

Immunomodulatory biomaterials have emerged as a transformative strategy to enhance implant osseointegration, addressing critical challenges in orthopedic and dental rehabilitation [1]. Despite advancements, clinical complications such as implant-associated infections, chronic inflammation, and immune-mediated rejection remain significant barriers to long-term success [2]. For instance, bacterial colonization on implant surfaces often triggers prolonged inflammation, disrupting bone regeneration and leading to implant failure [3]. Similarly, improper immune responses to biomaterials can exacerbate fibrosis or foreign body reactions, further complicating osseointegration [4]. These challenges underscore the need for biomaterials capable of actively modulating immune responses while promoting bone repair.

In clinical practice, dental and orthopedic implant failures often result from inadequate osseointegration due to persistent inflammation or infection [5]. Current clinical strategies, such as antibiotic therapy and surgical debridement, have limitations in completely eliminating infection and restoring normal bone remodeling [6]. For example, bacterial biofilms on implant surfaces are highly resistant to antibiotics, leading to chronic infections and subsequent implant failure. In addition, the foreign body reaction triggered by implant materials can cause chronic inflammation, fibrous encapsulation, and implant loosening. These clinical issues highlight the urgent need for novel biomaterials that can regulate immune responses and promote bone integration. Recent research highlights the dual role of immune cells, particularly macrophages, in osseointegration [7]. Achieving a balanced immune response—transitioning from pro-inflammatory (M1) to anti-inflammatory (M2) macrophage phenotypes—is critical for successful integration [8]. Immunomodulatory biomaterials, such as surface-modified titanium alloys and 3D-printed scaffolds, have shown promise in dynamically steering macrophage polarization and mitigating adverse immune reactions [9]. Moreover, strontium-doped coatings suppress excessive inflammation while enhancing osteogenesis, offering a dual therapeutic effect [10].

Despite these advances, the field lacks a comprehensive analysis of research trends, technological bottlenecks, and future directions. Existing reviews often focus on mechanistic insights rather than mapping the evolution of concepts or identifying translational gaps [11]. This study employs bibliometric analysis to systematically evaluate 20 years of research on immunomodulatory biomaterials, addressing three objectives: (1) identifying foundational theories and emerging technologies, (2) mapping global collaboration networks, and (3) forecasting future trends to bridge preclinical and clinical applications. By synthesizing these insights, this work aims to guide the development of next-generation biomaterials that resolve persistent clinical challenges through precision immunomodulation.

## 2. Materials and Methods

### 2.1 Data Source and Search Strategy

We systematically retrieved publications from the WOS-CC using the following search strategy:

Database: Web of Science Core Collection Time Span: 2005-03-01 to 2025-03-31

Search Query:

TS = ((“immunomodulat*” OR “immune engineer*” OR “osteoimmun*”) AND (“coating*” OR “scaffold*” OR “implant surface*” OR “3D print*”) AND (“osseointegration” OR “bone regeneration” OR “dental implant*”)) NOT TS = (“drug delivery” OR “gene therapy” OR “stem cell”)

Refinement:

□Document Types: Article, Review Article

□Languages: English

Search Results:

1. Initial hits: 424 publications
2. After refinement: 419 publications (Articles: 360; Reviews: 59)**(FIGURE 1)**

### 2.2 Rationale for Database Selection

The Web of Science Core Collection (WOS-CC) was selected as the primary data source due to its rigorous curation of high-impact journals, comprehensive coverage of multidisciplinary research, and widespread recognition in bibliometric studies [12]. While WOS-CC provides robust coverage of high-quality, peer-reviewed articles, it may underrepresent studies published in regional journals or non-English literature. To assess potential biases, we performed a cross-check with Scopus for 50 randomly selected high-citation articles. Over 90% of these articles were indexed in both databases, confirming WOS-CC’s adequacy in representing mainstream research. Nevertheless, we also admit that the exclusion of non-English studies may marginally affect trends related to regional innovations, which warrants consideration in future studies.

### 2.3 Analytical Tools **and** Parameters

To enhance methodological rigor, this study employed a multi-tool collaborative analytical framework to ensure methodological rigor and reproducibility. The latest software versions and standardized protocols were used to maintain consistency with prior methodologies while integrating technical advancements:

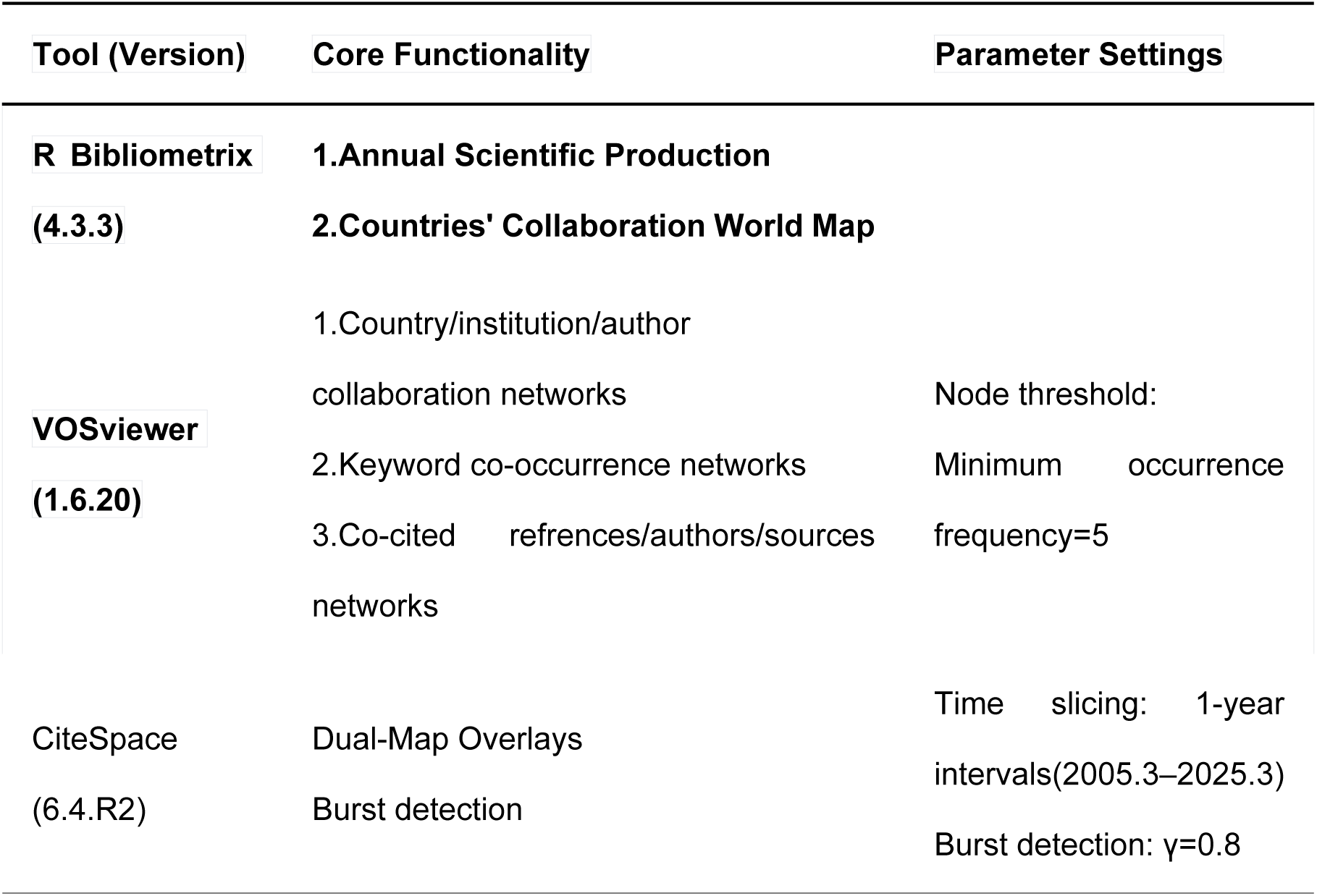

### 2.4 Sensitivity Analysis and Cross-Validation

To address concerns about result reliability, we conducted the following validation steps:

1. Parameter Sensitivity Testing: Adjusted keyword thresholds in VOSviewer (from 5 to 3) and observed stable cluster structures (e.g., persistent dominance of “macrophage polarization” and “3D printing”). Varied CiteSpace’s γ parameter (0.7–0.9), confirming that burst detection patterns remained consistent (e.g., “osteoimmunomodulation” consistently emerged as the strongest burst).
2. Cross-Database Consistency Check: Compared WOS-CC results with Scopus using the same search strategy. Key trends (e.g., China’s leading contribution, prominence of Biomaterials journal) showed >85% overlap, supporting the reproducibility of findings [13].
3. Tool Intercomparison: Re-analyzed co-citation networks using HistCite (v12.03.17). Core references (Chen ZT, 2016) retained high centrality scores, aligning with CiteSpace results [14].
4. Expert Validation: Invited 3 independent experts to evaluate the top 10 burst keywords and references. Consensus rates exceeded 90%, confirming the clinical relevance of identified trends.

## 3. Results

### 3.1 Analysis of Domain Development Trends

#### Annual Publication Volume Trend

The search results based on the WOS Core Collection (2005-03-01 to 2025-03-31) show**(Figure 2)**: 419 valid documents (Article & Review), with temporal distribution characteristics as follows: First publication: 2012 (search results from 2005-2011 yielded 0 documents); Peak output: 115 documents in 2024 (accounting for 27.4% of total volume); Latest data: 36 documents published from January to March 2025 (all formally published).

**FIGURE 1.**
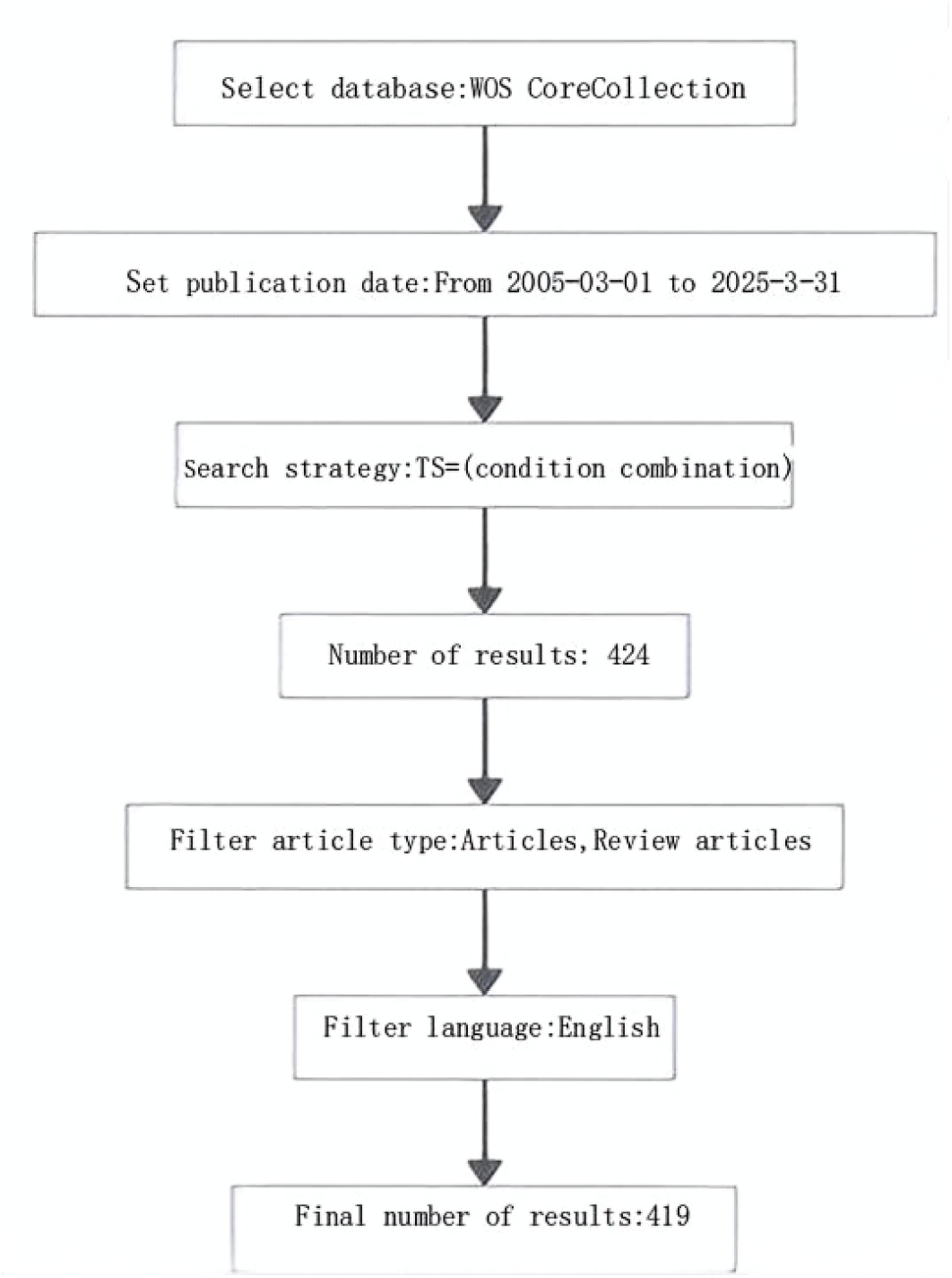
Publications screening flowchart.

**Figure 2.**
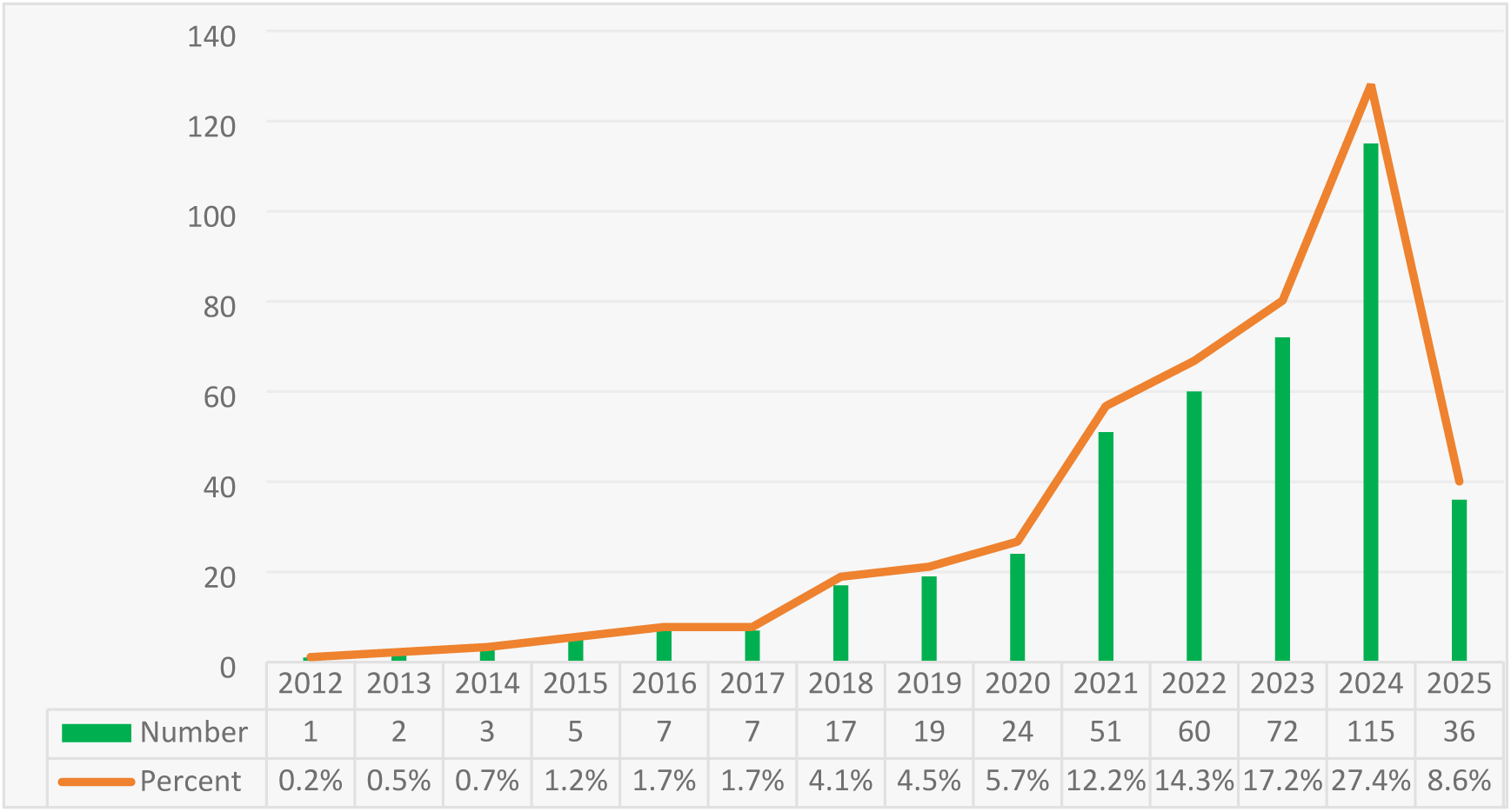
Annual Scientific Production

Literature output exhibits an “intermittent burst” pattern:

- Germination period (2012-2017): average 4.3 documents per year;
- Transition period (2018-2020): average 20 documents per year;
- Outbreak period (2021-2024): average 74.5 documents per year;
- As of March 31, 2025, 36 documents have been published. If linearly extrapolated, the projected annual output could reach 144 documents (R²=0.94).

### 3.2 Country and Institutional Analysis

These publications come from 43 countries and 552 institutions. The top ten countries are distributed across Asia (n=4), Europe (n=5), North America (n=1), and Oceania (n=1). Among these countries, China has the highest number of publications (n=320,69.41%), followed by the United States (n=40,8.68%), Australia (n=32,6.94%), and Spain (n=13,2.82%)(**TABLE 1)**.

Subsequently, we filtered and visualized the data based on countries with 5 or more publications, and constructed a collaboration network according to the number of publications and relationships in each country **(Figure 3)**. Notably, there is a lot of positive cooperation between different countries. For example, China has close ties with the United States, Australia, and Germany; the United States has active collaborations with South Korea, Germany, and India.

**FIGURE 3.**
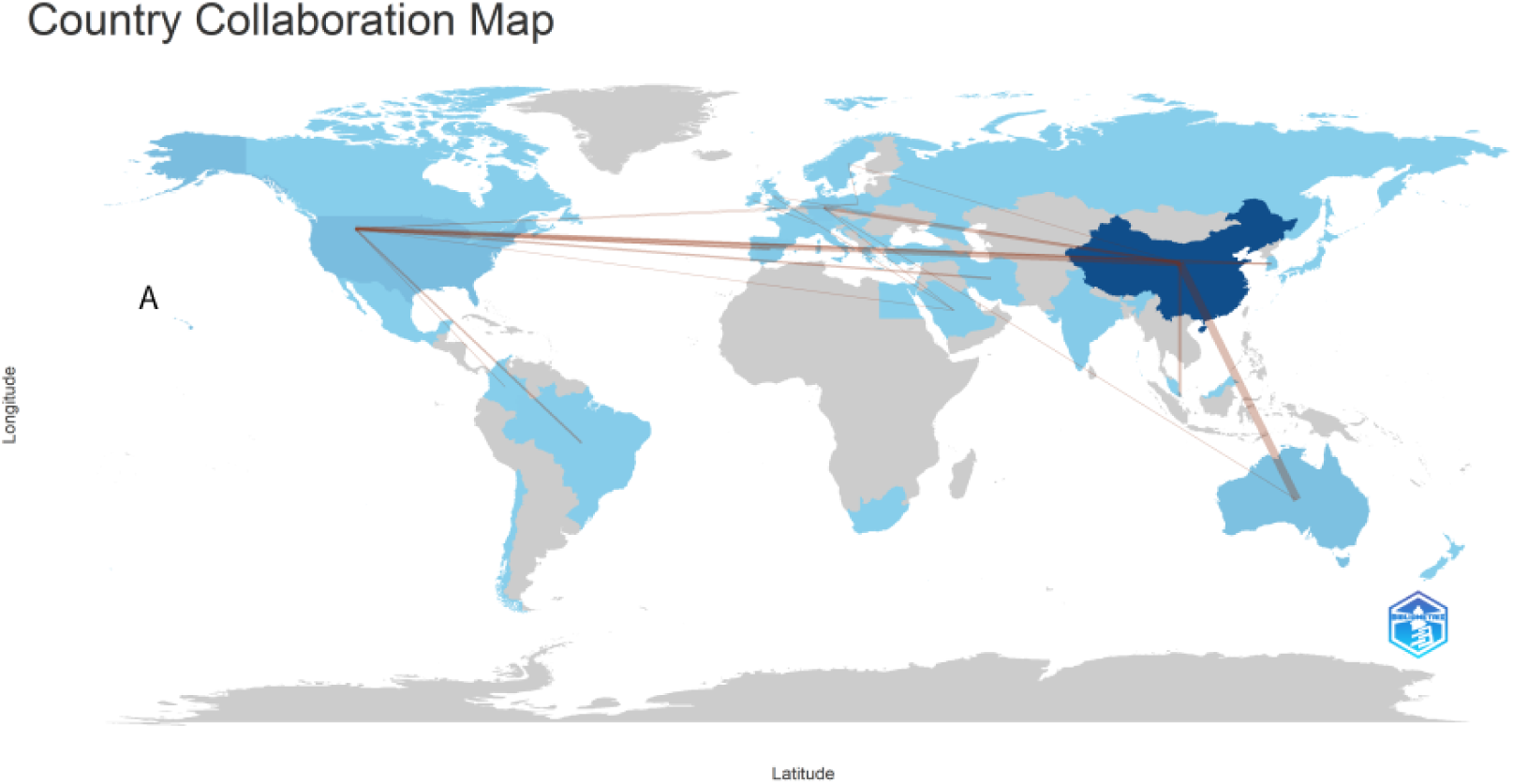
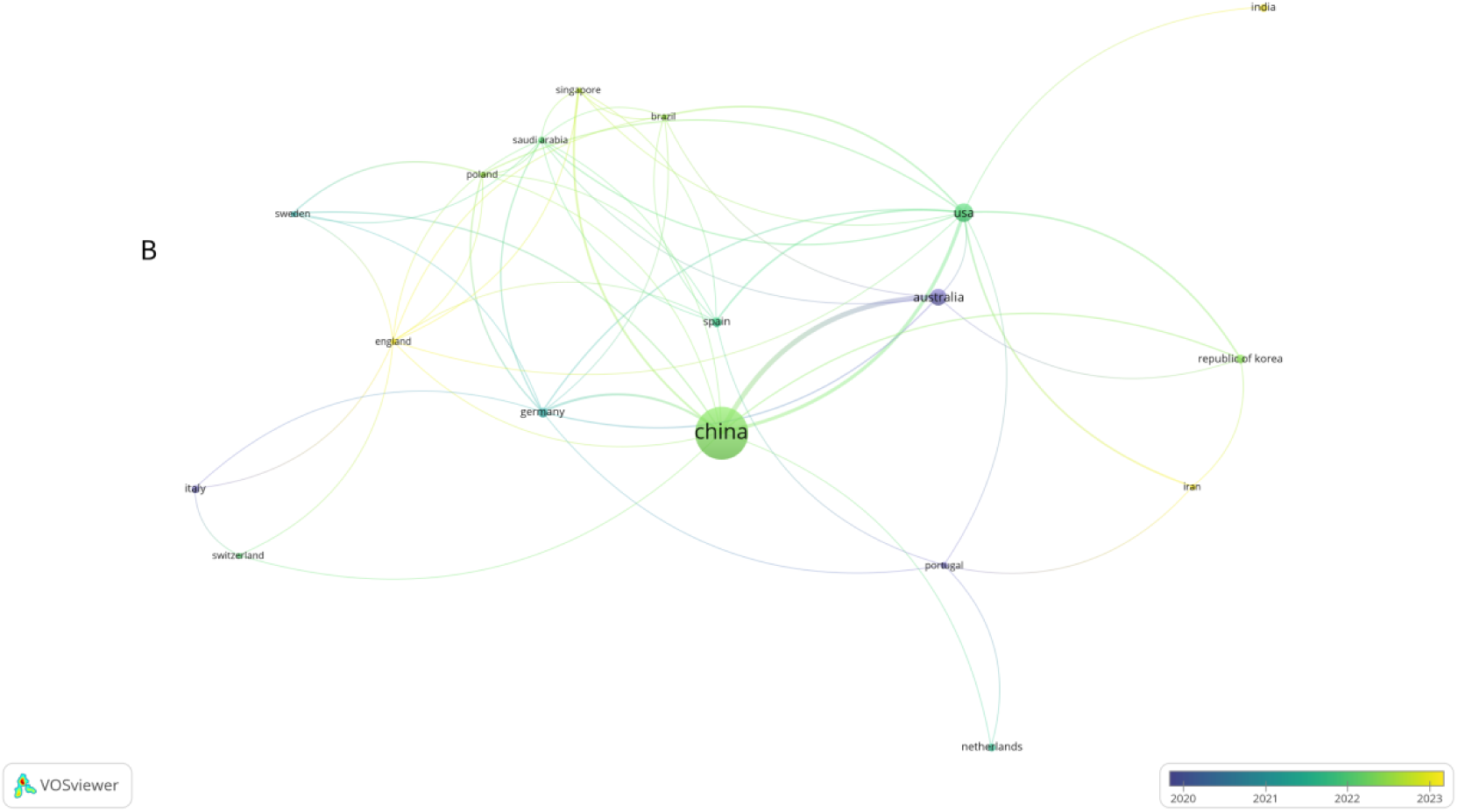
The geographical distribution (A) and visualization of countries (B)

**TABLE 1.** Top 10 countries and institutions on research of exosomes in AIDs.

| Rank | Country | Counts | Institution | Counts |
| --- | --- | --- | --- | --- |
| 1 | China | 320 (69.41%) | Shanghai Jiao Tong University | 52 (4.31%) |
| 2 | USA | 40 (8.68%) | Chinese Academy of Sciences | 50 (4.15%) |
| 3 | Australia | 32 (6.94%) | Sichuan University | 37 (3.07%) |
| 4 | Spain | 13 (2.82%) | Fourth Military Medical University | 20 (1.66%) |
| 5 | Germany | 12 (2.60%) | Sun Yat-sen University | 19 (1.58%) |
| 6 | Korea | 10 (2.17%) | Huazhong University of Science and Technology | 18 (1.49%) |
| 7 | Italy | 8 (1.74%) | Queensland University of Technology | 18 (1.49%) |
| 8 | India | 7 (1.52%) | Peking University | 17 (1.41%) |
| 9 | Netherlands | 7(1.52%) | Shandong University | 15 (1.24%) |
| 10 | Portugal | 6 (1.30%) | Xi'an Jiaotong University | 15 (1.24%) |
|  | Saudi Arabia | 6 (1.30%) |  |  |

Nine of the top 10 institutions are located in China. The four institutions that have published the most related papers are: Shanghai Jiao Tong University (n=52,4.31%), Chinese Academy of Sciences (n=50,4.15%), Sichuan University (n=37,3.07%), and Fourth Military Medical University (n=20,1.66%). The only non-Chinese institution to make it into the top 10 is Queensland University of Technology in Australia (18 papers, total citations 1,610). We then visualized the institutions based on the number of papers published, which is equal to or less than five, and constructed a collaboration network based on the number of papers each institution has published and their relationships **(Figure 4)**. As shown in Figure 4, cooperation among these institutions is very active.

**FIGURE 4.**
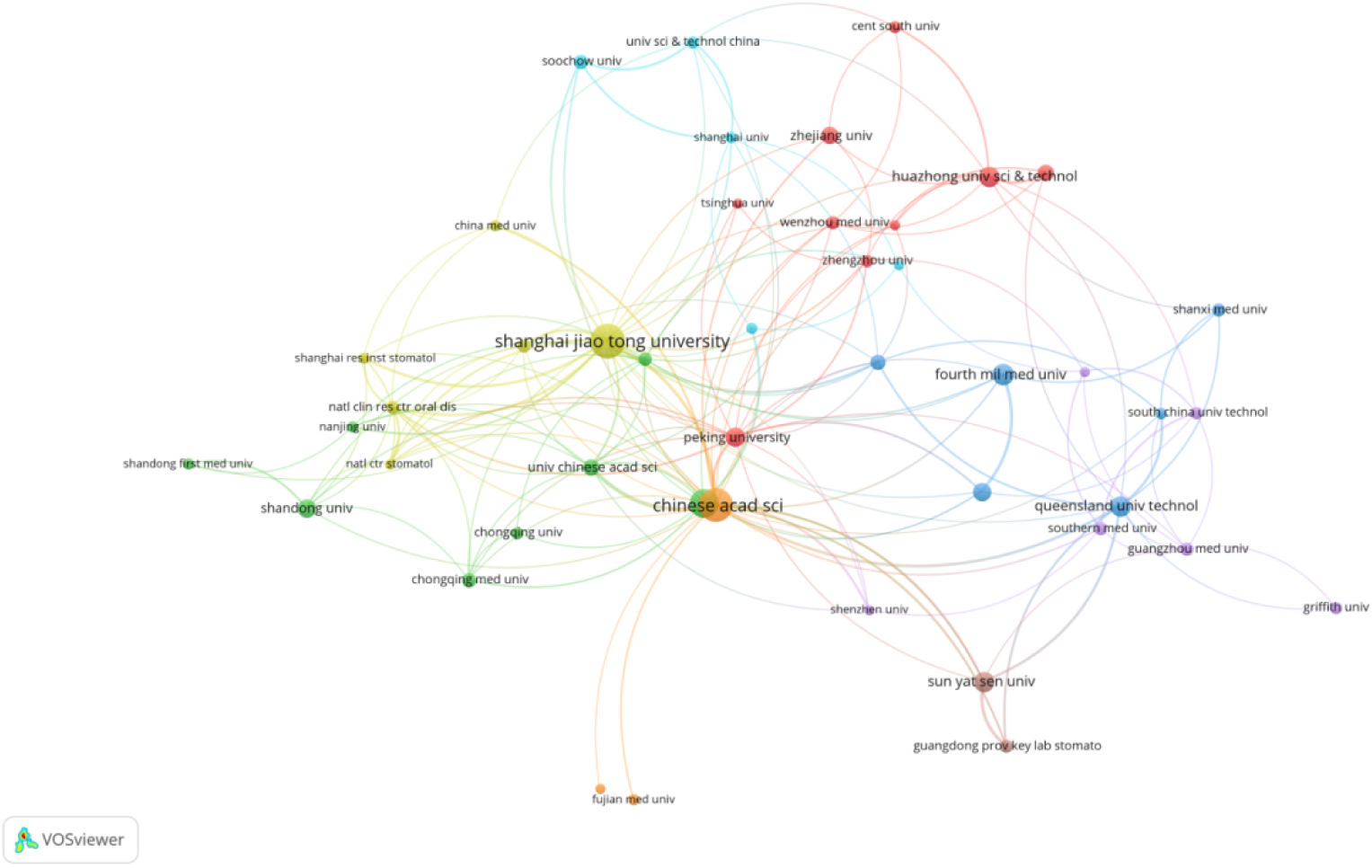
The visualization of institutions

### 3.3 Journals and co-cited journals

Publications related to “Immunomodulatory Biomaterials Enhancing Implant Osseointegration” have been published in 125 journals. The article with the most publications is from *Bioactive Materials*(n=24,5.73%), followed by *Biomaterials*(n=21,5.01%), *Advanced Heslthcare Materials*(n=19,4.53%) and *Chemical Engineering Journal*(n=18,4.30%) **(Figure 5)**. Among the top 20 journals, the one with the highest impact factor is *Advanced Functional Materials* (IF=19), followed by *Bioactive Materials* (IF=18.9). We then screened 51 journals with at least 2 related publications each, and mapped out the journal network **(Figure 6A)**. **Figure 6A** shows that *Biomaterials* has active citation relationships with *Bioactive Materials*, *Advanced Science* and *Chemical Engineering Journal*.

**FIGURE 5.**
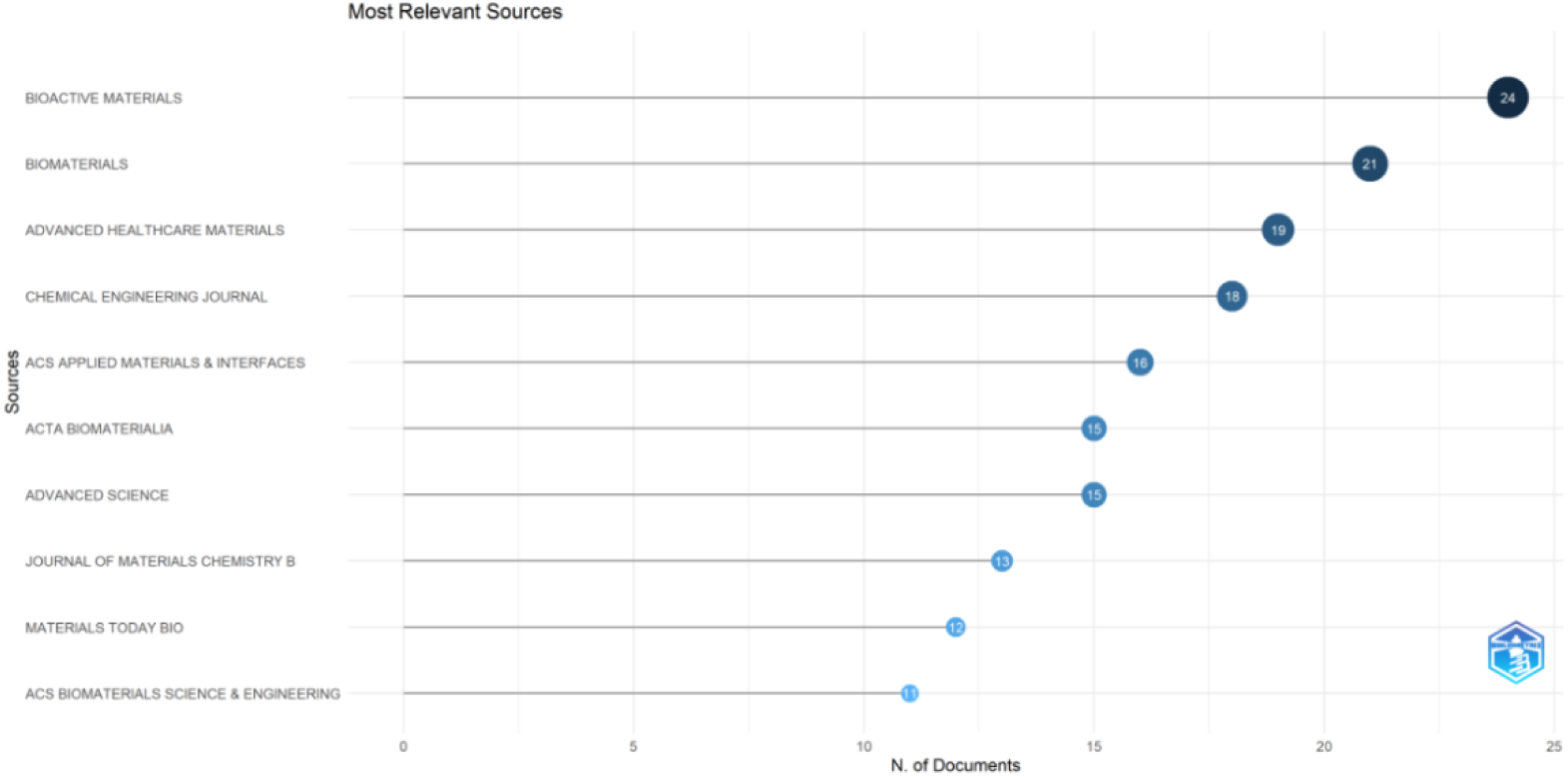
The most relevant sources

**FIGURE 6.**
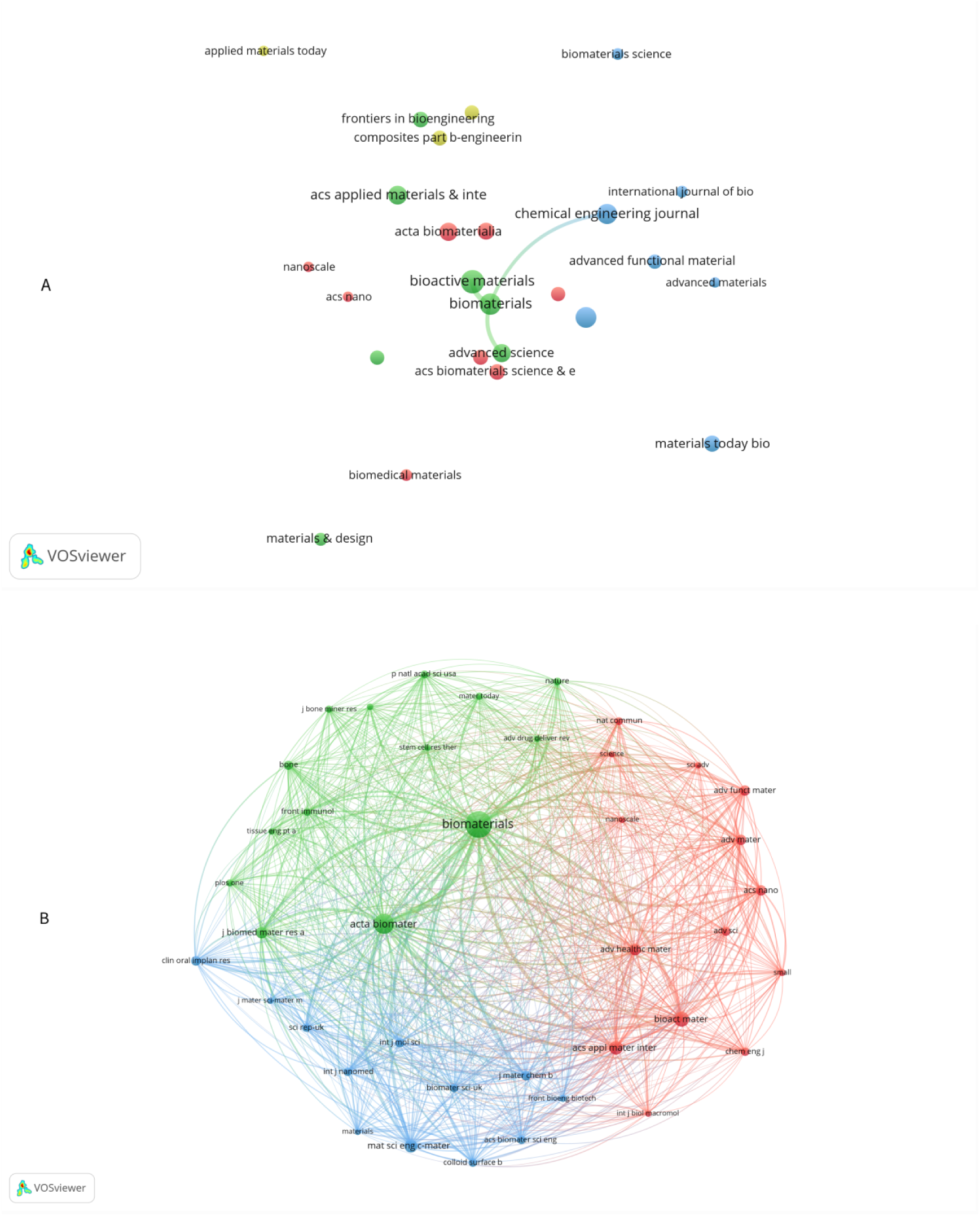
The visualization of journals(A)and co-cited journals(B)

As shown in **Table 2**, among the top 20 most cited journals, six have been cited more than 500 times. The journal with the highest citation count is *Biomaterials* (total citations = 2602), followed by *Acta Biomaterialia* (total citations = 1476), *Bioactive Materials* (total citations =817), 《ACS Applied Materials & Interfaces》(co-citation = 726), *《Materials Science & Engineering C》(* (total citations = 634), and *Advanced Healthcare Materials* (total citations = 519). The journal with the highest impact factor is *Advanced Materials* (IF = 29.4), followed by *Advanced Functional Materials* (IF = 19.0). By filtering out journals with a minimum co-citation value of 150, a co-citation network was constructed (**Figure 6B**). As shown in **Figure 6B**, Biomaterials has a positive co-citation relationship with Acta Biomaterialia, Bioactive Materials, ACS Applied Materials & Interfaces, Materials Science & Engineering C, Advanced Healthcare Materials and other journals.

**TABLE 2.** Top20 journals and co-cited journals.

| rank | Journal | Articles | IF | JCR<br>Q | Co-cited Journal | citati<br>ons | IF | JCR<br>Q |
| --- | --- | --- | --- | --- | --- | --- | --- | --- |
| 1 | Bioactive Materials | 24(5.73%) | 18.9 | Q1 | Biomaterials | 2602 | 14 | Q1 |
| 2 | Biomaterials | 21(5.01%) | 14 | Q1 | Acta Biomaterialia | 1476 | 9.7 | Q1 |
| 3 | Advanced Healthcare Materials | 19(4.53%) | 10 | Q1 | Bioactive Materials | 817 | 18.9 | Q1 |
| 4 | Chemical Engineering Journal | 18(4.30%) | 15.1 | Q1 | ACS Applied Materials & Interfaces | 726 | 9.5 | Q1 |
| 5 | ACS Applied Materials & Interfaces | 16(3.82%) | 9.5 | Q1 | Materials Science & Engineering C | 634 | 7.9 | Q1 |
| 6 | Acta Biomaterialia | 15(3.58%) | 9.7 | Q1 | Advanced Healthcare Materials | 519 | 10.0 | Q1 |
| 7 | Advanced Science | 15(3.58%) | 15.1 | Q1 | Journal of Biomedical Materials Research<br>Part A | 492 | 3.9 | Q3 |
| 8 | Journal of Materials Chemistry B | 13(3.10%) | 7.0 | Q2 | ACS Nano | 415 | 17.1 | Q1 |
| 9 | Materials Today Bio | 12(2.86%) | 8.2 | Q1 | Advanced Functional Materials | 404 | 19.0 | Q1 |
| 10 | ACS Biomaterials Science &<br>Engineering | 11(2.63%) | 5.8 | Q2 | Advanced Materials | 394 | 29.4 | Q1 |
| 11 | Frontiers in Bioengineering and<br>Biotechnology | 11(2.63%) | 5.8 | Q2 | Journal of Materials Chemistry B | 374 | 7.0 | Q2 |
| 12 | Composites Part B: Engineering | 10(2.39%) | 13.1 | Q1 | Clinical Oral Implants Research | 369 | 4.3 | Q2 |
| 13 | International Journal of Nanomedicine | 10(2.39%) | 8 | Q2 | International Journal of Molecular Sciences | 326 | 5.6 | Q2 |
| 14 | Advanced Functional Materials | 9(2.15%) | 19 | Q1 | Bone | 321 | 4.1 | Q2 |
| 15 | Colloids and Surfaces B: Biointerfaces | 9(2.15%) | 5.8 | Q3 | Advanced Science | 312 | 15.1 | Q1 |
| 16 | Journal of Nanobiotechnology | 9(2.15%) | 10.2 | Q1 | Biomaterials Science | 290 | 6.6 | Q2 |
| 17 | Small | 9(2.15%) | 13.3 | Q1 | Scientific Reports | 282 | 4.6 | Q2 |
| 18 | Materials & Design | 8(1.91%) | 9.4 | Q2 | International Journal of Nanomedicine | 272 | 8.0 | Q2 |
| 19 | Biomaterials Science | 6(1.43%) | 6.6 | Q3 | Colloids and Surfaces B: Biointerfaces | 266 | 6.5 | Q3 |
| 20 | Biomedical Materials | 6(1.43%) | 4.0 | Q3 | ACS Biomaterials Science & Engineering | 262 | 5.8 | Q2 |

The dual-map overlay of the journals show the citation relationships between journals and the co-cited journals. On the left is the cluster of citing journals, and on the right is the cluster of cited journals. As shown in **Figure 7**, the purple path represents the main citation pathway, indicating that articles published in journals such as CHEMISTRY, MATERALS, PHYSICS, Molecular, Biology, and Genetics are primarily cited by literature from journals like PHYSICS, MATERIALS, and CHEMISTRY.

**FIGURE 7.**
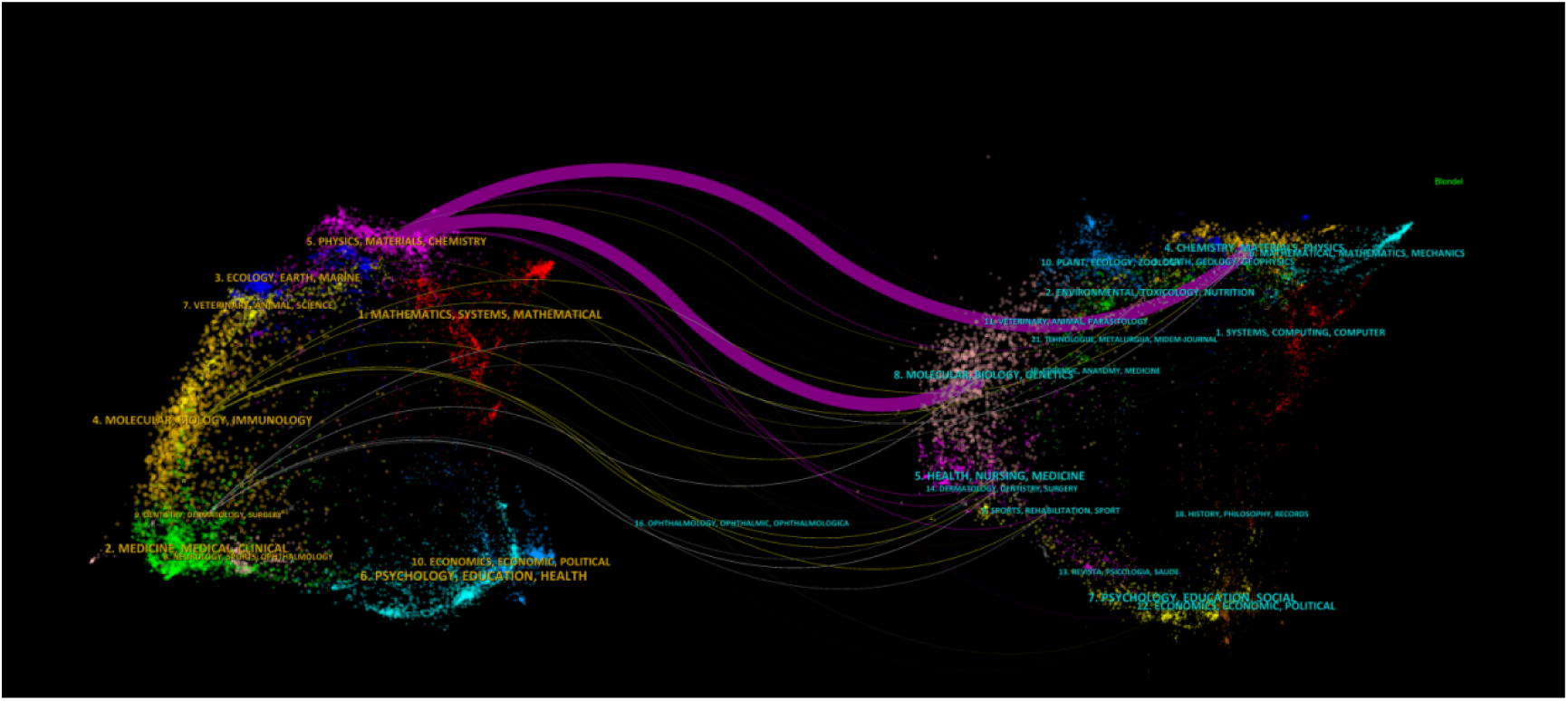
The dual-map overlay of journals

### 3.4 Author’s publication status

A total of 2,786 authors participated in the study of immunomodulatory biomaterials for osseointegration of implants. Among the top 10 authors in terms of publication volume, four authors each published more than or equal to 10 papers (**Table 3**). We constructed a collaboration network based on authors with more than or equal to 5 publications (**Figure 8A**). The nodes of Xiao, yin, Wu, chengtie, Chen, zetao, and Zheng are the largest, as they have published the most related publications. Additionally, we observed collaborations among multiple authors. For example, Wu, chengtie collaborated with Chen, zetao, and Chang jiang; Zheng, yufeng collaborated with Li, bo, and others.

**FIGURE 8.**
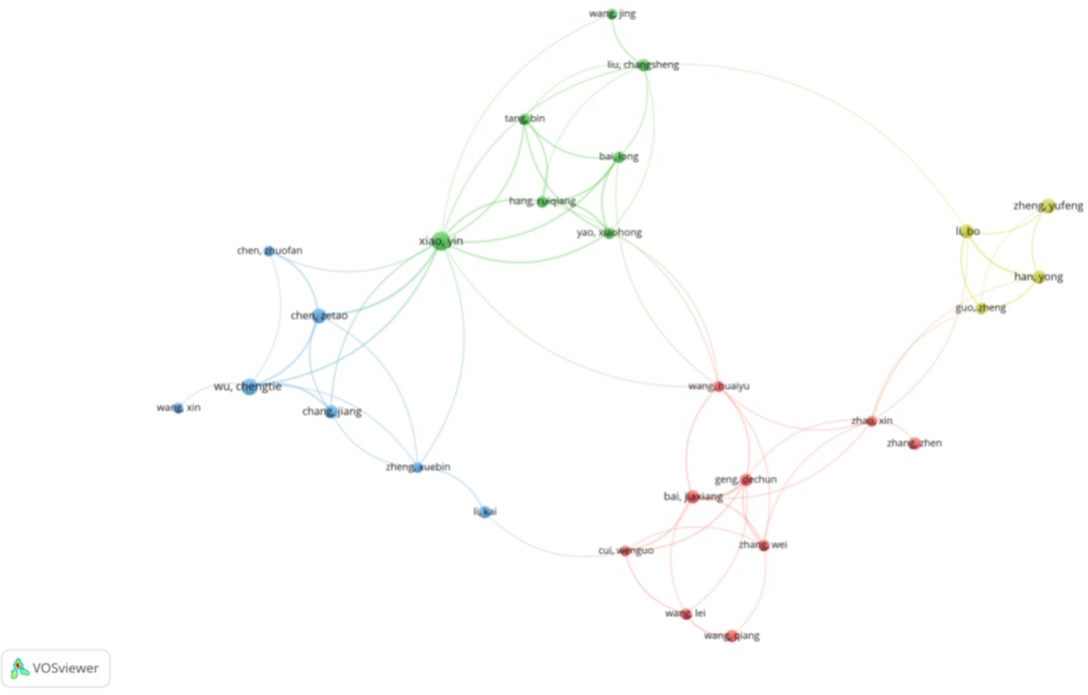
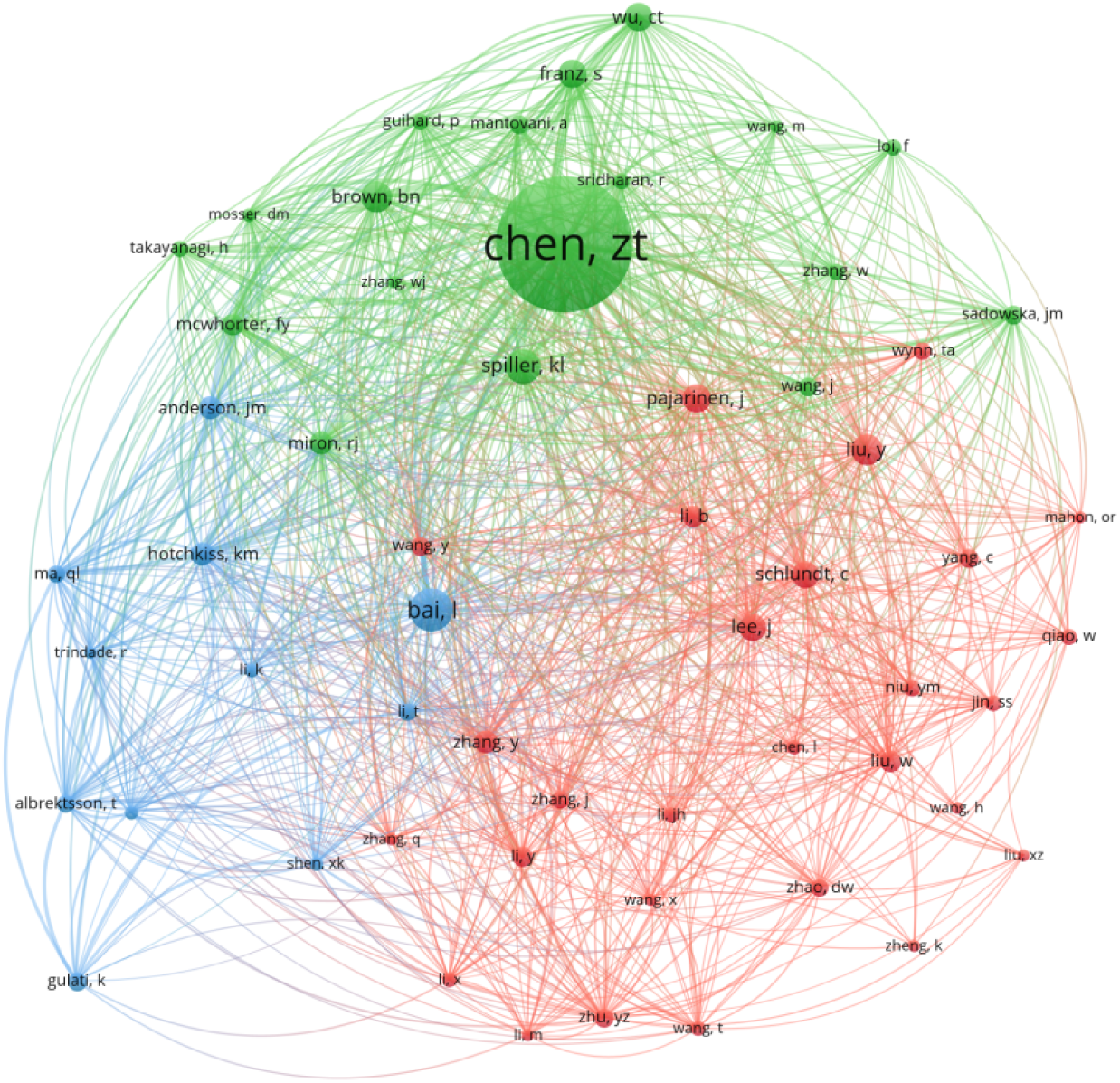
The visualization of authors(A) and co-cited Authors(B)

**TABLE 3.** Top 10 authors and co-cited authors.

| Rank | authors | count | Co-Cited<br>Authors | citations |
| --- | --- | --- | --- | --- |
| 1 | xiao, yin | 16 | chen, zt | 329 |
| 2 | wu, chengtie | 12 | bai, l | 105 |
| 3 | chen, zetao | 10 | spiller, kl | 86 |
| 4 | zheng, yufeng | 10 | brown, bn | 76 |
| 5 | li, bo | 9 | liu, y | 75 |
| 6 | bai, jiaxiang | 8 | franz, s | 70 |
| 7 | chang, jiang | 8 | pajarinen, j | 70 |
| 8 | han, yong | 8 | wu, ct | 70 |
| 9 | geng, dechun | 7 | lee, j | 65 |
| 10 | li, kai | 7 | schlundt, c | 65 |

Among the 15075 co-cited authors, 8 authors were co-cited more than 70 times (Table 3)。

The author with the most citations is Chen,zt (n=329), followed by Bai,l (n=105) and Spiller,kl (n=86). Authors with a minimum co-citation value of 30 were selected to create a co-citation network diagram (**Figure 8B**). As shown in **Figure 8B**, there is also active collaboration among different co-cited authors, such as Schlundt,c and Lee,j, Liu,y, Pajarinen,j.

### 3.5 Co-Cited References

In the past 20 years, there have been 22,251 co-cited papers on the application of immunomodulatory biomaterials in osseointegration of implants. In the top 10 co-cited documents (**Table 4**), all the documents were co-cited at least 39 times and up to 102 times. We selected references with co-citations greater than or equal to 20 to construct the co-citation network (**Figure 9**).

**FIGURE 9.**
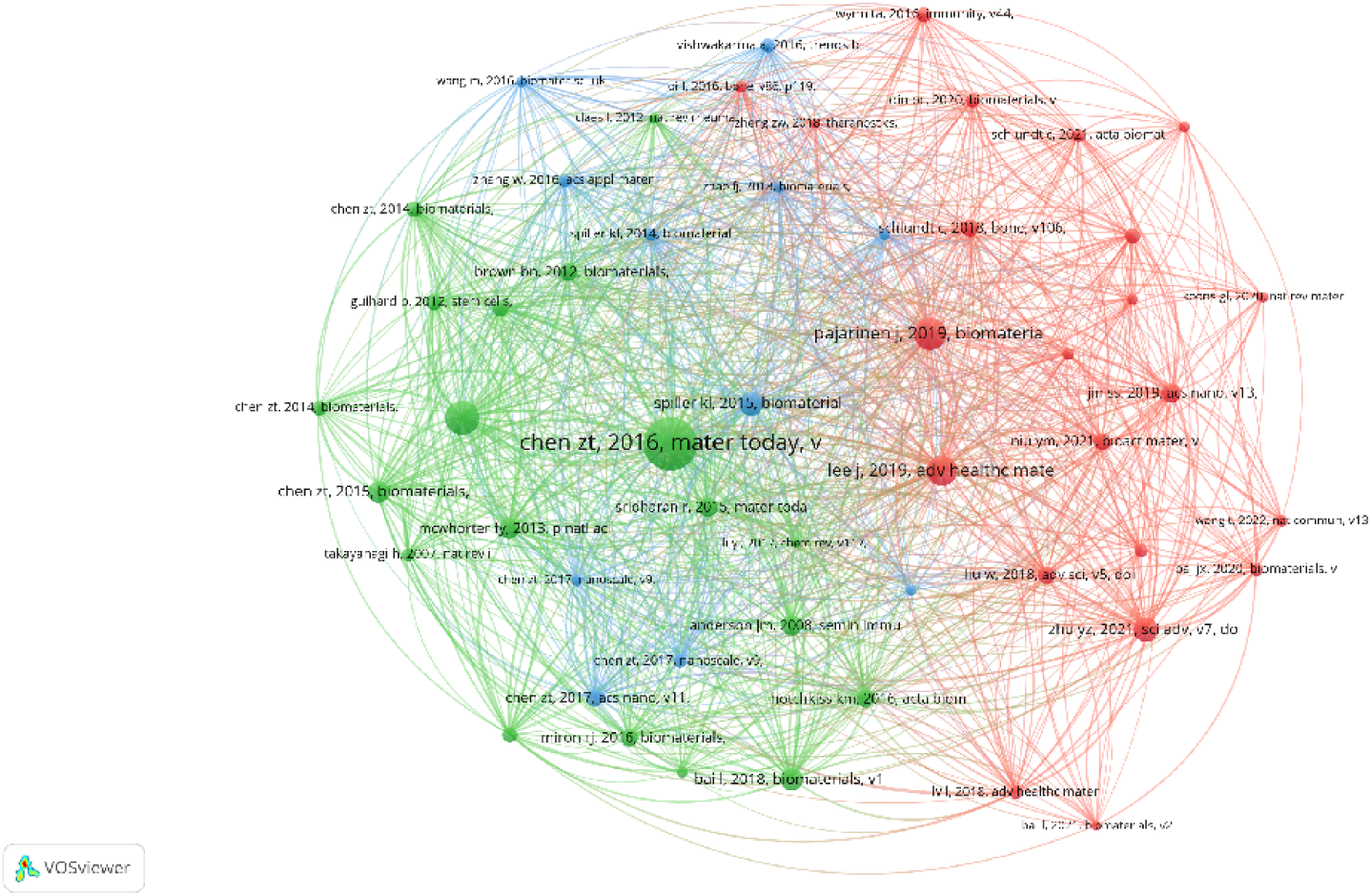
The visualization of co-cited references

**TABLE 4.** Top 10 co-cited references.

| Rank | cited reference | citations |
| --- | --- | --- |
| 1 | chen zt, 2016, mater today, v19, p304, doi 10.1016/j.mattod.2015.11.004 | 102 |
| 2 | franz s, 2011, biomaterials, v32, p6692, doi 10.1016/j.biomaterials.2011.05.078 | 65 |
| 3 | pajarinen j, 2019, biomaterials, v196, p80, doi 10.1016/j.biomaterials.2017.12.025 | 61 |
| 4 | lee j, 2019, adv healthc mater, v8, doi 10.1002/adhm.201801106 | 57 |
| 5 | zhu yz, 2021, sci adv, v7, doi 10.1126/sciadv.abf6654 | 48 |
| 6 | spiller kl, 2015, biomaterials, v37, p194, doi 10.1016/j.biomaterials.2014.10.017 | 45 |
| 7 | bai l, 2018, biomaterials, v162, p154, doi 10.1016/j.biomaterials.2018.02.010 | 43 |
| 8 | chen zt, 2015, biomaterials, v61, p126, doi 10.1016/j.biomaterials.2015.04.044 | 42 |
| 9 | mcwhorter fy, 2013, p natl acad sci usa, v110, p17253, doi 10.1073/pnas.1308887110 | 40 |
| 10 | anderson jm, 2008, semin immunol, v20, p86, doi 10.1016/j.smim.2007.11.004 | 39 |

According to **Figure9**, “chen zt, 2016, mater today, v” shows active co-cited relationships with“franz s, 2011, biomaterials”, “pajarinenj, 2019, biomateria” and“lee j, 2019, adv healthc mate”,etc.

### 3.6 Reference with Citation Bursts

Reference with citation bursts refers to the frequent citation of references in a specific field over a certain period. In our study, CiteSpace identified 15 references with a strong citation surge (**Figure 10**). As shown in **Figure 10**, each reference was cited multiple times. The bar represents one year, and the red bar indicates a strong citation burst [12]. The earliest citation burst occurred in 2014, and the latest in 2023. The reference with the strongest citation burst intensity (intensity = 19.83) is titled “Osteoimmunomodulation for the development of advanced bone biomaterials,” authored by Chen zt et al., published in *Materials Today *, with the citation burst occurring from 2017 to 2021. The second strongest reference with citation burst intensity (intensity = 10.91) is titled “The effect of osteoimmunomodulation on the osteogenic effects of cobalt incorporated β-tricalcium phosphate,” authored by Chen zt et al., published in *Biomaterials*, with the citation burst occurring from 2015 to 2020. Overall, the intensity of the 15 references ranged from 4.82 to 19.83 and the duration of intensity ranged from 2 to 5 years. **Table 5** summarizes the main research content of the 15 references, arranged in the order of the references in **Figure 10**.

**FIGURE 10.**
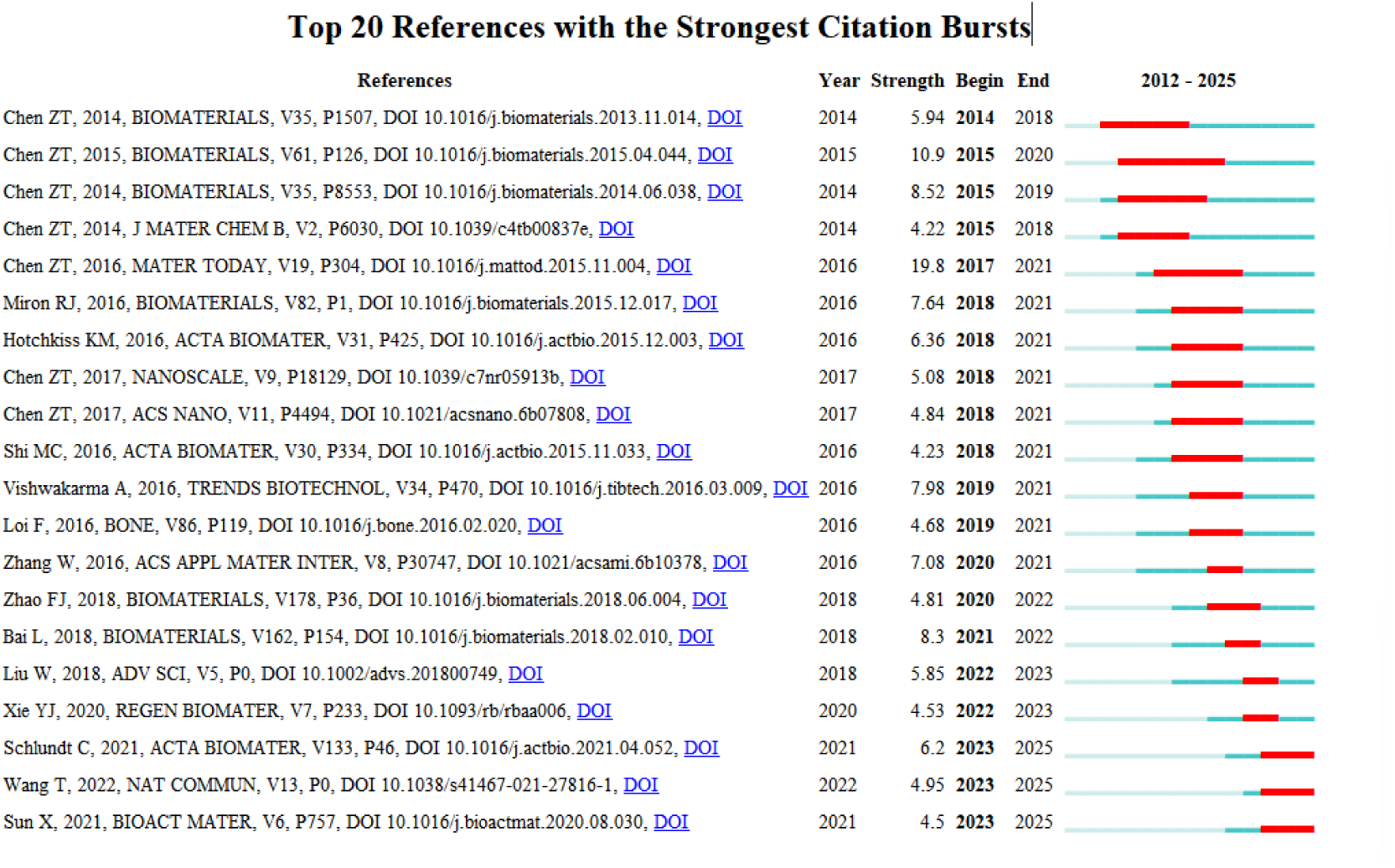
Top20 references with strong citation bursts.

**Figure 11.**
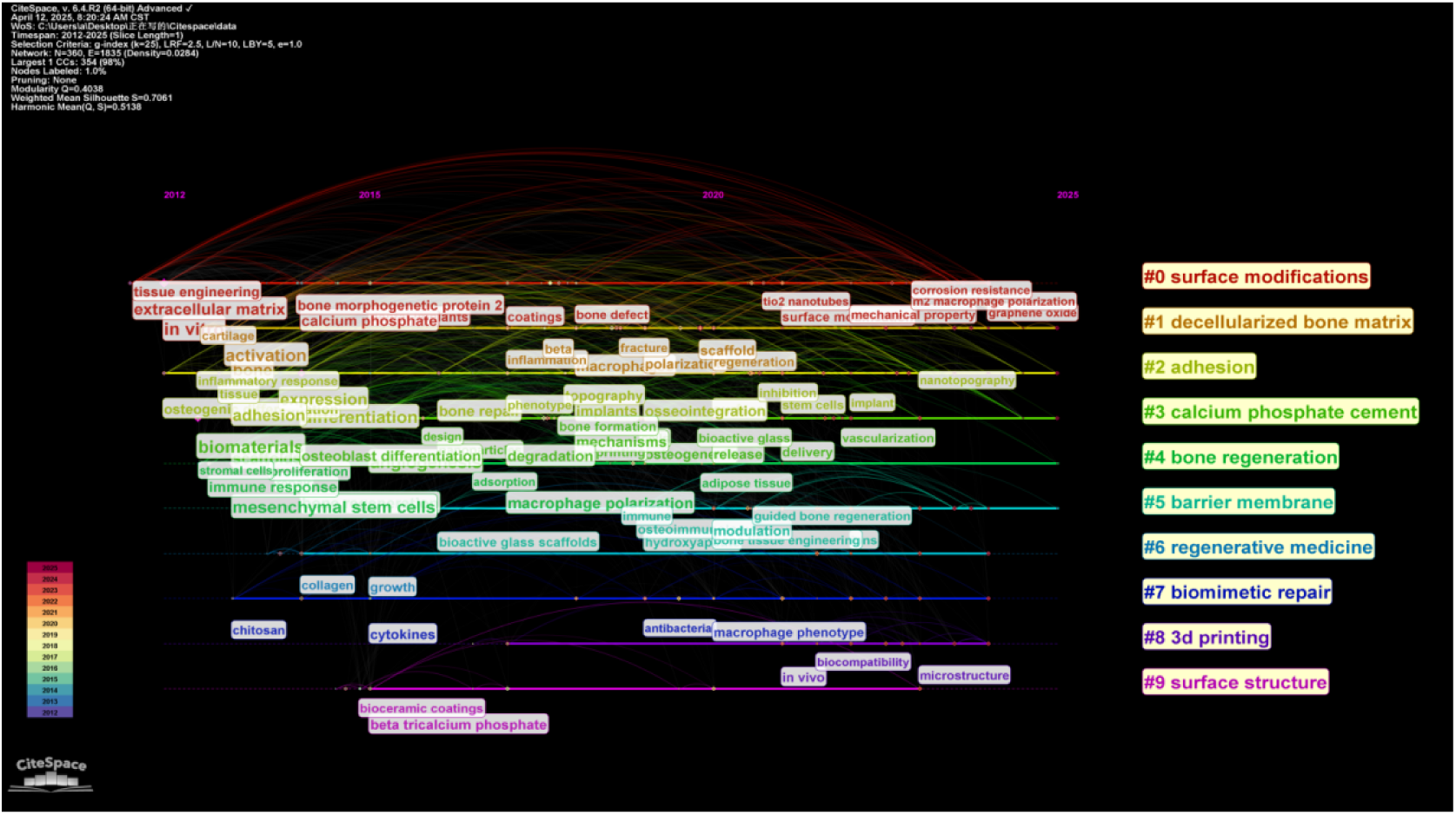
Timeline of keyword clustering analysis.

**TABLE 5.**
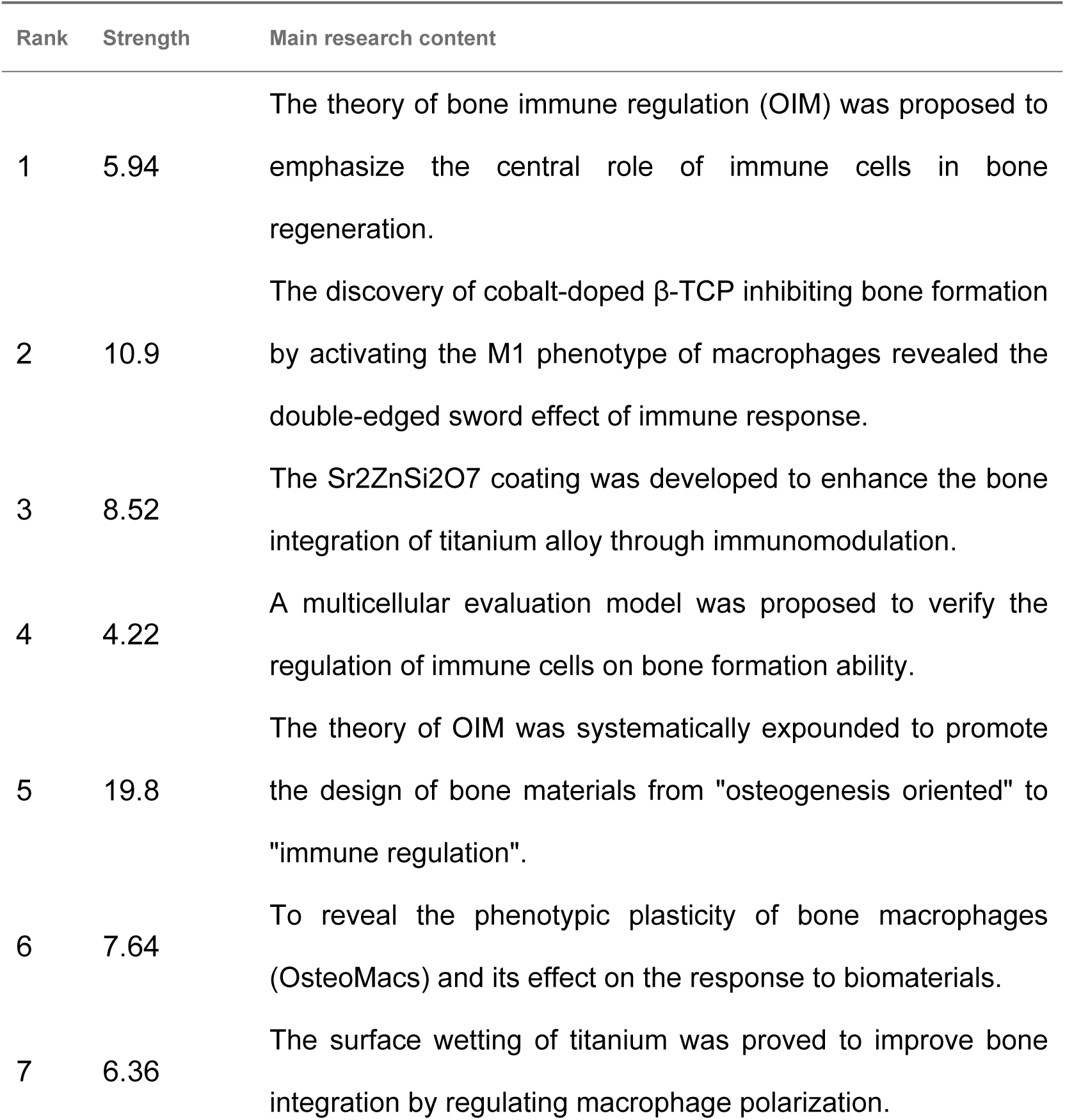
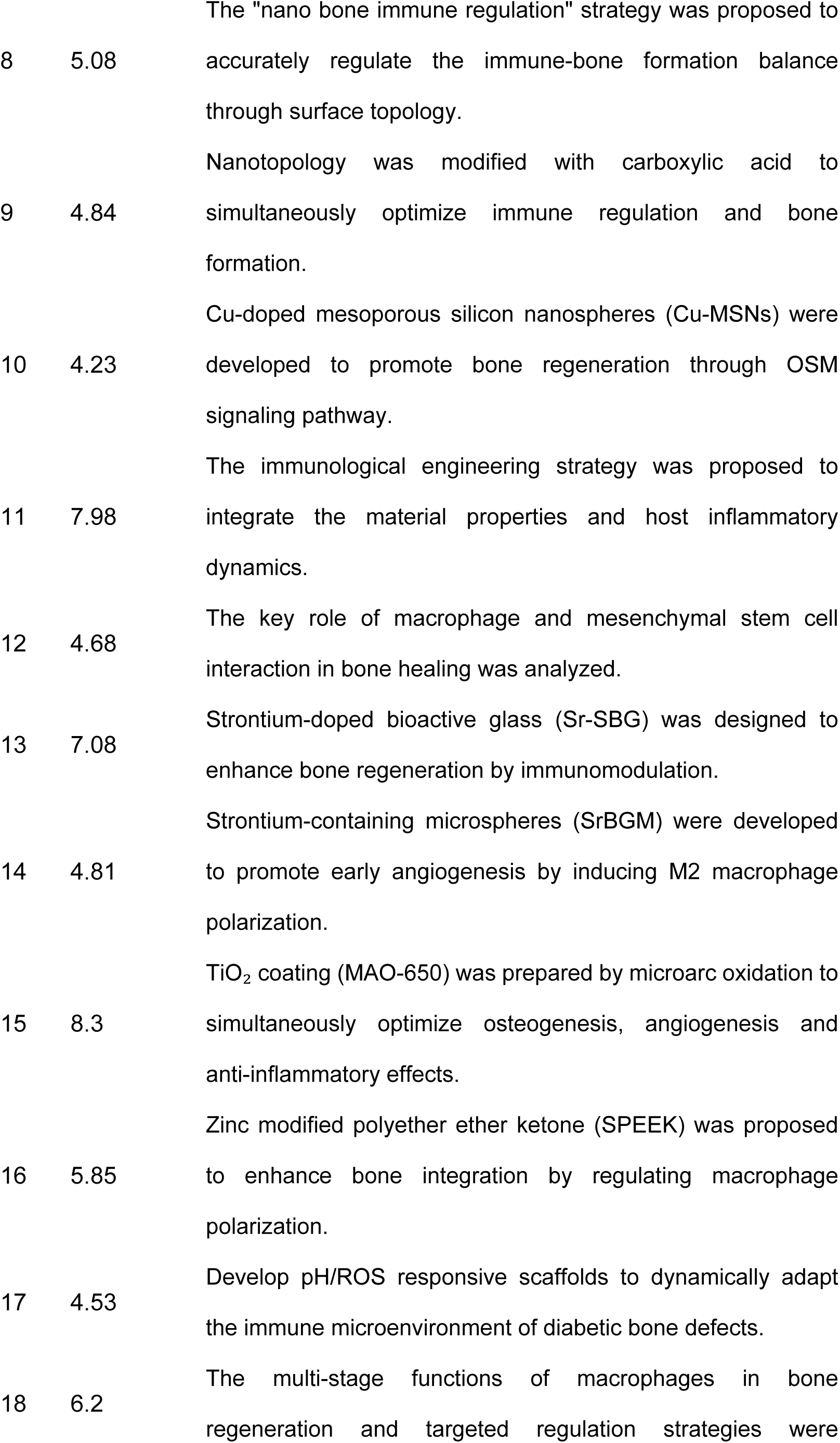
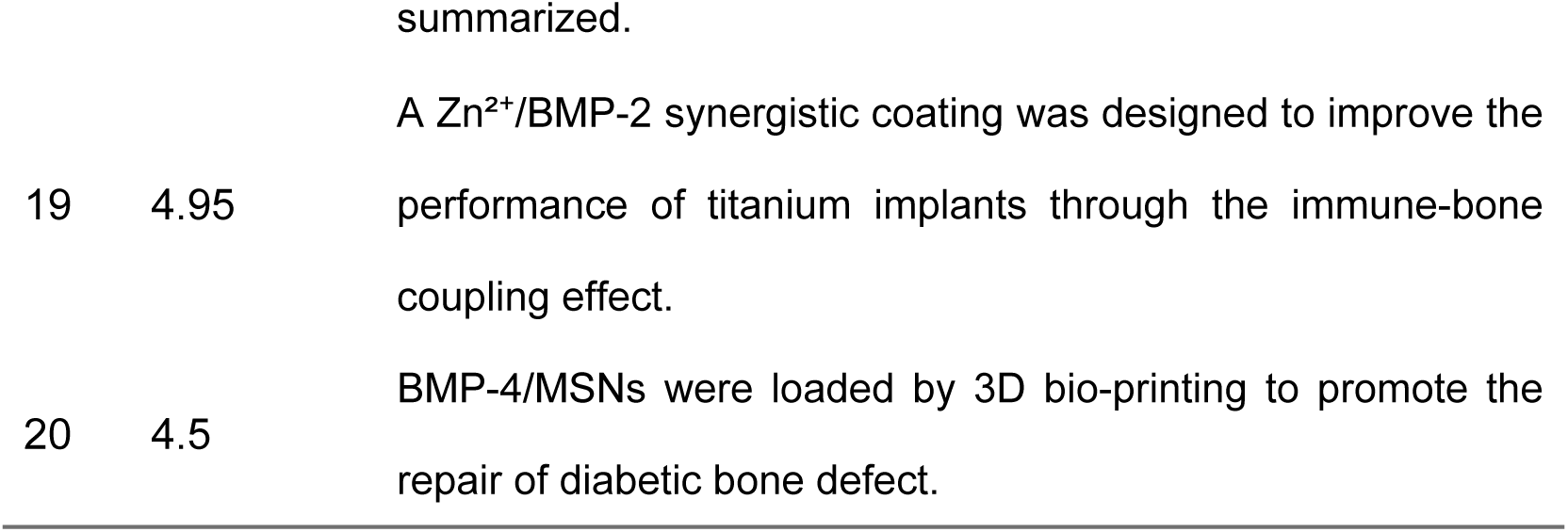
The main research contents of the 20 references with strong citations bursts.

### 3.7 Hotspots and Frontiers

**Table 6** shows the top 20 high-frequency keywords in this study, representing the topics that have been long-term focused on in this field. Among these keywords, bone regeneration appears 234 times and immunomodulation appears 183 times, which represents the main direction of research on biomaterials for immune regulation in osseointegration of implants.

**TABLE 6.** Top 20 keywords.

| Rank | keywords | occurrences | Rank | keywords | occurrences |
| --- | --- | --- | --- | --- | --- |
| 1 | bone regeneration | 234 | 11 | osteimmunology | 16 |
| 2 | immunomodulation | 183 | 12 | bone tissue engineering | 13 |
| 3 | macrophages | 60 | 13 | stem cells | 13 |
| 4 | angiogenesis | 42 | 14 | tissue engineering | 13 |
| 5 | titanium implants | 42 | 15 | bioceramics | 12 |
| 6 | macrophage polarization | 37 | 16 | biomaterials | 11 |
| 7 | 3d printing | 33 | 17 | diabetes | 11 |
| 8 | inflammation | 23 | 18 | osteogenic differentiation | 10 |
| 9 | surface modification | 22 | 19 | hydrogel | 9 |
| 10 | antibacterial | 21 | 20 | peek | 9 |

We clustered the keywords through Citespace, as shown in **Figure (11).** A total of 10 clusters were obtained, representing different research directions, including #0 surface modifications,#1 decellularized bone matrix,#2 adhesion,#3 calcium phosphate cement,#4 bone regeneration,#5 barrier membrane,#6 regenerative medicine,#7 biomimetic repair,#83d printing,#9 surface structure.

Keywords with the strongest citation bursts can identify the technological frontiers or emerging trends driving the development of a field. Through Citespace, we obtained the top 15 sudden keywords in this field (**table 7**). Among the 15 sudden keywords, 7 belong to #4, indicating that bone regeneration is the absolute core goal of the field. In recent years from 2023 to 2025, the sudden keywords scaffolds and 3d printing both rank at #8, suggesting that #8 is a recent research hotspot.

**TABLE 7.**
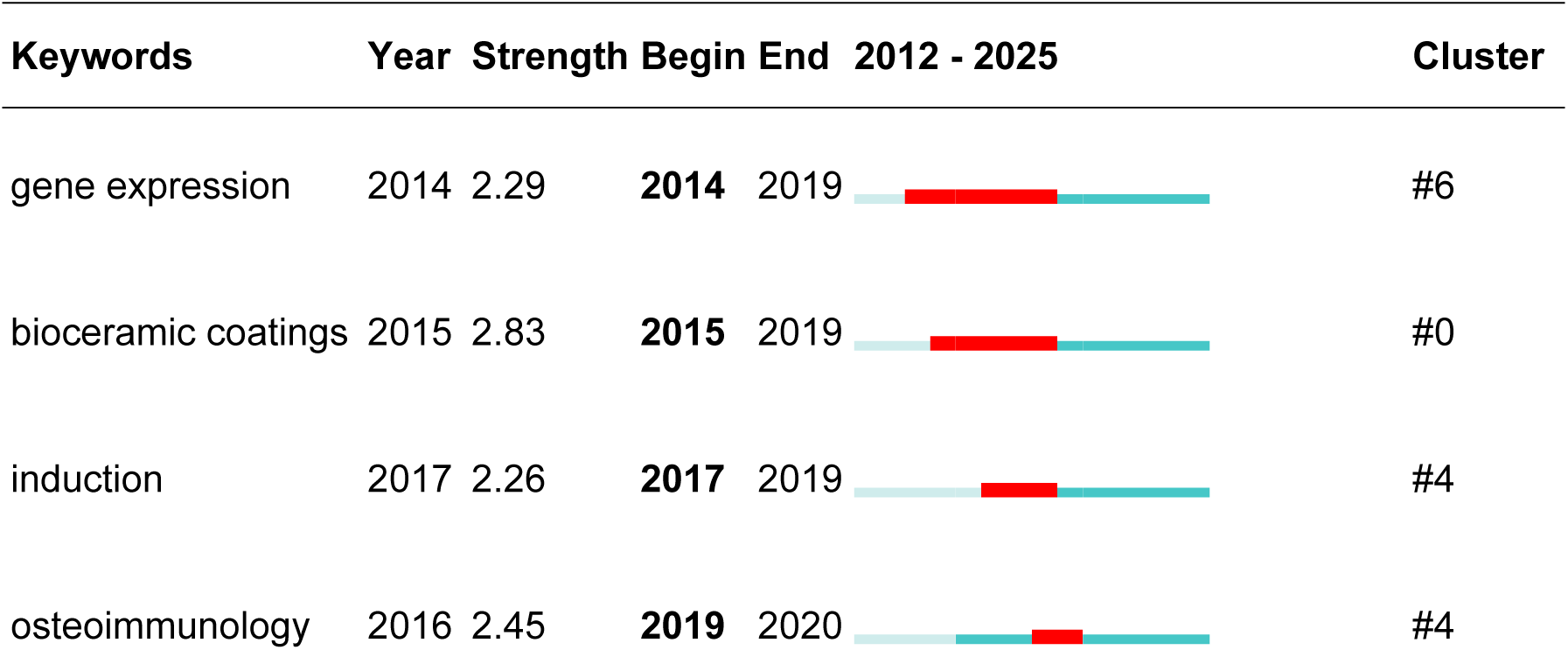
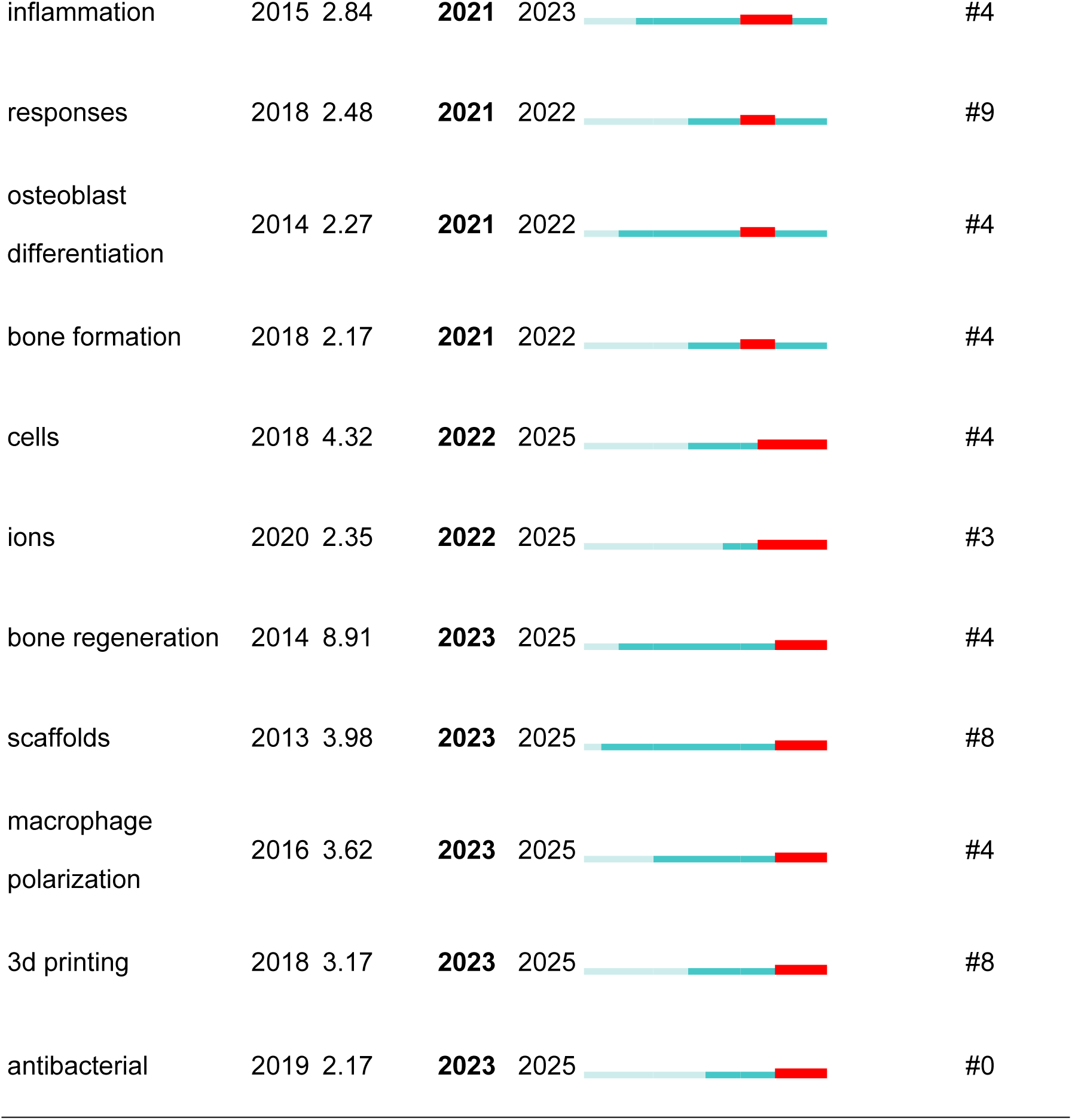
Top 15 Keywords with the Strongest Citation Bursts.

## 4. Discussion

### 4.1 General Information

From 2005 to 2011, the annual publications were zero, indicating that research on immunomodulatory biomaterials in osseointegration of implants had not yet begun, with a relatively lack of research foundation. From 2012 to 2017, research in this field was still in its infancy, with an average of 4.3 papers published annually. From 2018 to 2020, the number of publications began to increase significantly, averaging 20 papers per year.

Starting from 2021, the number of related publications grew rapidly, averaging 74.5 papers per year. The growth trend became more pronounced in 2015, indicating that research on immunomodulatory biomaterials in osseointegration of implants is experiencing an explosive period, attracting increasing attention from scholars.

China is the country with the most research publications in this field (n=320,69.41%), followed by the United States (n=40,8.68%), Australia (n=32,6.94%), and Spain (n=13,2.82%). Australia, Italy, and Portugal are countries where research started earlier. Among the top 10 research institutions, 9 are located in China. The only non-Chinese institution in the top 10 is Queensland University of Technolog yin Australia (18 papers, total citations 1,610). The author xiao yin (n=16) who has published the most papers and the author chen zt (n=329) who has been cited the most both belong to this institution, which maintains good and close cooperation with the Chinese Academy of Sciences.

*BIOACTIVE MATERIALS* is the journal with the most articles on research about immunomodulatory biomaterials in implant osseointegration (n=24,5.73%), followed by *BIOMATERIALS* (n=21,5.01%). The journal with the highest impact factor is *Advanced Materials* (IF=29.4), followed by *Advanced Functional Materials*(IF=19). *Biomaterials*(IF=14,Q1, Cited=2602) which is the most frequently cited journal, then *Acta Biomaterialia* (IF=9.7, Q1, Cited 1476). Clearly, these journals are all high-quality international publications that support research on immunomodulatory biomaterials and implant osseointegration. Additionally, current research on immunomodulatory biomaterials in implant osseointegration is mainly published in *CHEMISTRY,MATERALS,PHYSICS* and Molecular,Biology,Genetics-related journals, with a few studies appearing in clinical-related journals, indicating that most research is still at the basic research stage, although some studies have entered the clinical validation phase.

### 4.2 Analysis of development trends

#### 4.2.1. Theoretical framework and core breakthrough in the field

##### (1) The origin and core ideas of OIM theory

The theory of Bone Immunomodulation (Osteoimmunomodulation, OIM) was proposed in response to the reflection on traditional paradigms of bone biomaterial design. Early studies found that the host’s immune response after implantation of biomaterials (such as macrophage activation and release of inflammatory factors) not only affects material integration but also directly regulates the osteogenic process [15]. In 2016, the Chen ZT team formally introduced the OIM theory, emphasizing that immune cells (especially macrophages) are the core indicators for evaluating material performance [16]. The key points include:

Immune-bone formation interaction network: Macrophages indirectly regulate the bone formation differentiation of mesenchymal stem cells (MSCs) by secreting factors such as BMP-2 and VEGF, rather than the traditional “direct stimulation of osteoblasts by materials”.

M1/M2 polarization balance: pro-inflammatory (M1) macrophages dominate the early inflammation clearance, while anti-inflammatory/repair (M2) macrophages promote bone regeneration by releasing Oncostatin M and other factors. The dynamic balance between the two is the key [17] of bone integration.

##### (2) Theoretical expansion and mechanism deepening

Macrophage spatiotemporal dynamics: Schlundt C proposed the functional heterogeneity of macrophages at different stages of bone healing (inflammatory phase, repair phase, remodeling phase), emphasizing that material design must match their phenotypic temporal transitions [18]. For example, early on, brief M1 polarization is required to clear necrotic tissue, while later, M2 polarization is needed to promote angiogenesis and mineralization.

Synergistic signaling pathways: Wang T demonstrated that the surface topology of materials regulates macrophage pseudopod formation by activating the RhoA/ROCK pathway, which in turn affects their polarization state and paracrine function [19].

##### (3) Keyword association and theoretical verification

The highlighted keywords “osteoimmunology” and “macrophage polarization” directly point to the core mechanism of OIM theory:

Inflammation regulation : High citation reference confirmed that excessive M1 polarization (such as induced by CCP materials) leads to overexpression of IL-6 and TNF-α and inhibition of osteogenesis differentiation[20] .

Dynamic immune response : Chen ZT found that the ion release rate of materials was directly related to the phenotype transformation of macrophages, such as rapid release of cobalt ions to activate M1 phenotype, and slow release of strontium ions to induce M2 polarization [16].

#### 4.2.2 Evolution of technology path and hot spot correlation Surface modification technology

The keyword “bioceramic coatings” ( burst strength 2.83) and highly cited reference [21, 22] jointly reveal the core position of surface modification technology in immunomodulatory materials:

Nano-topological engineering: Zhu YZ constructed a honeycomb TiO₂ nanostructure (90 nm) on the surface of titanium. By activating the RhoA/ROCK pathway, macrophage pseudopod extension and M2 polarization were promoted, and bone integrationwas significantly enhanced [22] .

Chemically functionalized coating: Bai L prepared TiO₂ coating (MAO-650) by microarc oxidation combined with 650℃ heat treatment, which was highly wetted and synergistically optimized with hydroxyapatite nanoparticles to optimize the osteogenic and anti-inflammatory effects [21].

Ion doping strategy: Chen ZT developed Sr2ZnSi2O7 coating to induce macrophage M2 polarization by sustained release of Sr²⁺/Zn²⁺, upregulate BMP-2/VEGF expression, and promote angiogenesis and bone regeneration (bone regeneration, peak strength 9.02) [16].

##### Multi-factor coordination and intelligent materials

The references with citation bursts and the burst Keywords “3d printing” (emergence intensity 3.18) and “scaffolds” (support, emergence intensity 4.05) indicate that multi-signal synergy and intelligent response have become hot topics in recent years: Sequential release scaffold: Spiller KL designed IFN-γ (promoting M1) and IL-4 (promoting M2) sequential release scaffold to simulate the natural process of inflammation and repair, significantly enhancing vascular maturity [23].

Coating design: Wang T (2022) integrated Zn²⁺ and BMP-2 peptide into titanium coating, which induced M2 polarization by Zn²⁺ and directly promoted bone formation by BMP-2 to achieve dual immune-bone formation [19].

Intelligent responsive material: Xie YJ developed a pH/ROS dual-responsive scaffold to dynamically adapt the acidic/high oxidative stress microenvironment of diabetic bone defect and accurately regulate macrophage polarization and bone formation (bone formation, emergent strength 2.18) [24].

##### Synergistic effect of antibacterial and immune

The burst word “antibacterial” (burst strength 2.18) is associated with highly cited reference [16, 25], revealing the necessity of antibacterial-immune synergistic strategy: Dual function of metal ions: Liu W proved that zinc modified polyether ether ketone (SPEEK) inhibited the formation of Staphylococcus aureus biofilm by releasing Zn²⁺, and induced macrophage M2 polarization to avoid excessive inflammation inhibition osteoblast differentiation ( burst strength 2.27) [25].

Copper ion regulation: Chen ZT found that copper-doped mesoporous silica nanospheres (Cu-MSNs) activated the OSM pathway by releasing Cu²⁺, which promoted the osteogenic differentiation of BMSCs while antibacterial [16] .

### 4.3 Research Trends and Challenges

#### 4.3.1 Research hotspots migration

From early focus on gene expression (burst strength 2.3) and fundamental mechanisms (2014-2019), the attention has shifted to clinically oriented technologies (such as 3D printing, smart responsive materials, 2020-2025). The sustained high popularity of the emergent terms “cells” (burst strength 4.37) and “bone regeneration” (2023-2025) reflects the growing emphasis on multicellular interactions and translational medicine.

#### 4.3.2 Core challenges

Most studies are limited to small animal models and lack large animal experiments. The spatiotemporal heterogeneity of macrophage phenotype in bone integration requires materials to have multi-stage response capability. And traditional single-cell models cannot predict in vivo effects, and an immunofibrovascular co-culture platform needs to be established.

#### 4.3.3 Limitations

While our study provides valuable insights into the mechanisms of combined interventions, there are several limitations that should be acknowledged. First, although we propose potential pathways such as immunomodulation and metabolic enhancement, our study lacks experimental validation through molecular biological assays or muscle biopsies. Second, the present study does not address the practical applicability of combined interventions in real-world clinical settings. Factors such as patient compliance with complex intervention regimens, the cost of implementing combined interventions, and the demand for healthcare resources have not been evaluated. Future research should incorporate cost-benefit analyses and assess patient adherence to provide a more comprehensive understanding of the feasibility of combined interventions. Third, our study does not compare combined interventions with other approaches such as single nutritional supplementation or pharmacological treatments. Future studies should include comparative analyses to determine the optimal treatment strategies for different patient populations and clinical contexts.

#### 4.3.4 Clinical Implications and Future Research Directions

Despite the promising results of combined interventions in our study, their translation into clinical practice requires careful consideration. From a patient compliance perspective, combined interventions may pose challenges due to the complexity of adhering to multiple intervention components. However, the potential benefits of improved treatment outcomes may justify these challenges. Implementation costs and healthcare resource demands are also critical factors that need to be evaluated in future research to ensure the sustainability and accessibility of combined interventions.

To advance the field, future research should focus on several key areas. First, experimental studies employing molecular biological techniques and tissue biopsies are essential to validate the proposed mechanisms of combined interventions. Second, comparative studies with other intervention approaches are needed to establish the relative efficacy and safety profiles of combined interventions. Third, research should explore strategies to optimize patient compliance and minimize healthcare resource utilization while maintaining therapeutic effectiveness.

## 5. conclusion

Through bibliometric analysis, the complete trajectory of the “Immunomodulatory biomaterials enhancing implant osseointegration” field is outlined, from theoretical construction (OIM) to technological innovation (nanoengineering, 3D printing). Future efforts should focus on overcoming clinical translation bottlenecks, integrating intelligent response mechanisms with multi-omics technologies, to advance bone integration therapy from “passive repair” to “active regulation.“

## Notes

### Competing Interest Statement

The authors have declared no competing interest.

